# Maturation of cortical microstructure and cognitive development in childhood and adolescence: a T1w/T2w ratio MRI study

**DOI:** 10.1101/681221

**Authors:** Linn B. Norbom, Jaroslav Rokicki, Dag Alnæs, Tobias Kaufmann, Nhat Trung Doan, Ole A. Andreassen, Lars T. Westlye, Christian K. Tamnes

## Abstract

The restructuring and optimization of the cerebral cortex from early childhood and through adolescence is an essential feature of human brain development, underlying immense cognitive improvements. Beyond established morphometric cortical assessments, the T1w/T2w ratio quantifies partly separate biological processes, and might inform models of typical neurocognitive development and developmental psychopathology. In the present study, we computed vertex-wise T1w/T2w ratio across the cortical surface in 621 youths (3-21 years) sampled from the Pediatric Imaging, Neurocognition, and Genetics (PING) study and tested for associations with individual differences in age, sex, and both general and specific cognitive abilities. The results showed a near global linear age-related increase in T1w/T2w ratio across the brain surface, with a general posterior to anterior increasing gradient in association strength. Moreover, results indicated that boys in late adolescence had regionally higher T1w/T2w ratio as compared to girls. Across individuals, T1w/T2w ratio was negatively associated with general and several specific cognitive abilities mainly within anterior cortical regions. Our study indicates age-related differences in T1w/T2w ratio throughout childhood, adolescence and young adulthood, in line with the known protracted myelination of the cortex. Moreover, the study supports T1w/T2w ratio as a promising surrogate measure of individual differences in intracortical brain structure in neurodevelopment.

## Introduction

Childhood and adolescence is a time of extensive brain maturation and reorganization, shaped by multiple genetically based biological processes in dynamic interplay with the environment (Blakemore, 2012; Brown & Jernigan, 2012; Kremen et al., 2013; Lebel & Deoni, 2018). Magnetic resonance imaging (MRI) indicates that structural cortical development is characterized by widespread, but regionally variable and coordinated nonlinear decreases in volume and thickness, and early increases followed by smaller decreases in surface area (Amlien et al., 2016; Khundrakpam et al., 2019; Tamnes et al., 2017). This cortical maturation is mirrored by immense cognitive improvements from early childhood to young adulthood, with the development of basic functions such as sensorimotor processes preceding complex functions such as top-down control of behavior (Akshoomoff et al., 2014; Casey, Tottenham, Liston, & Durston, 2005). Detailed knowledge about cortical maturational patterns in youth is critical for models of normative neurocognitive development, and for serving as a blueprint for detecting neurodevelopmental deviations linked to maladjustment and psychopathology.

In addition to cortical morphometry, less explored measures of signal intensity variation in the T1-weighted (T1w) image may reflect additional and partly distinct neurobiological properties and processes (Andrews et al., 2017; Lewis, Evans, & Tohka, 2018; Lewis, Fonov, Collins, Evans, & Tohka, 2019; Norbom et al., 2019; Salat et al., 2009). Maturational and aging related cortical thinning have, for instance, been teased apart by their inverse relationships with cortical T1w intensity, which increases during development and declines in older aging (Westlye et al., 2010). More recently, T1w/T2w ratio has been suggested as a useful and possibly more specific measure of cortical microstructure (M. F. Glasser & Van Essen, 2011; Van Essen & Glasser, 2014). A handful of studies support the utility of T1w/T2w ratio for mapping global and regional maturational and aging related patterns (H. Grydeland et al., 2019; Håkon Grydeland, Walhovd, Tamnes, Westlye, & Fjell, 2013; Shafee, Buckner, & Fischl, 2015), and associations with cognitive performance in youth (Håkon Grydeland et al., 2013; Vandewouw, Young, Shroff, Taylor, & Sled, 2019) and adults (H. Grydeland, Westlye, Walhovd, & Fjell, 2016).

Although the underlying biology of the T1w/T2w ratio is controversial (Hagiwara et al., 2018; Ritchie, Pantazatos, & French, 2018; Uddin, Figley, Solar, Shatil, & Figley, 2019) and likely complex, the measure has been linked to differential myelination of the cerebral cortex (Ganzetti, Wenderoth, & Mantini, 2014; M. F. Glasser & Van Essen, 2011; Nakamura, Chen, Ontaneda, Fox, & Trapp, 2017). Intracortical myelination is a crucial feature of postnatal brain development, allowing for efficient signal transmission and structural support (Baumann & Pham-Dinh, 2001; Liu, Li, Zhu, Li, & Liu, 2019; Waxman & Bennett, 1972), and generally follows a posterior to anterior maturational pattern (Yakovlev & Lecours, 1967), with additional regional specificity (Deoni, Dean, Remer, Dirks, & O’Muircheartaigh, 2015; Nieuwenhuys, 2013; Ziegler et al., 2019). Cortical dysmyelination has also been linked to neurodevelopmental psychopathology (Bartzokis, 2005; Insel, 2010). Although T1w/T2w ratio shows promise for capturing distinct patterns of typical age-related differences in cortical microstructure across the lifespan, no previous study has tested the notion in typically developing children with an age range spanning below primary school years. Moreover, previous lifespan studies have included a restricted number of children or adolescents, consequently with limited statistical power to explore interactions between age and sex. Also, no previous study has tested the sensitivity of T1w/T2w ratio to general- and an array of specific cognitive abilities in typically developing youths.

To this end, we computed vertex-wise T1w/T2w ratio across the cortical surface in 621 youths (3-21 years) from the Pediatric Imaging, Neurocognition, and Genetics (PING) sample, and tested for associations between T1w/T2w ratio and age, sex and both general and specific cognitive abilities. We hypothesized that the T1w/T2w ratio would generally show a positive age association, putatively reflecting protracted intracortical myelination, but with regional variation. Next, given previous findings of limited sex differences in morphometric cortical development (Vijayakumar et al., 2016; Wierenga, Bos, van Rossenberg, & Crone, 2019), we predicted that boys and girls would show similar age related associations with T1w/T2w ratio. Finally, we tested for associations between T1w/T2w ratio and general and specific cognitive abilities, and predicted a positive association, based on the direction of reported developmental T1w/T2w findings (H. Grydeland et al., 2019; Shafee et al., 2015).

## Materials and Methods

### Participants

The publicly available PING study database (http://ping.chd.ucsd.edu) consists of data from a large sample of typically developing children and adolescents aged 3-21 years, with standardized behavioral measures, whole genome genotyping, and multimodal imaging for a large subgroup (Jernigan et al., 2016). Fluent English speakers within the desired age range were recruited across the US through local postings and outreach facilities. Exclusion criteria included somatic illness, history of head trauma, contraindications for MRI, preterm birth, diagnosis of mental disorders (except ADHD), mental retardation, and severe illicit drug use by mother during pregnancy. Subjects 18 years and above provided written informed consent, while written parental informed consent was obtained for all other subjects, in addition to child assent for subjects aged 7-17 (Jernigan et al., 2016).

From 998 subjects with available MRI and demographic data, T2-weighted (T2w) data was missing or acquired at insufficient resolution (> 1.2mm voxel size in any direction) for 322 subjects. Of the remainder, we additionally excluded 55 subjects after stringent quality control (see below), yielding a final sample of 621 subjects (303 girls) aged 3.2-21.0 years (mean = 12.4, SD = 4.8). Ten of these subjects had missing cognitive data and were excluded from relevant analyses. See Table 1 for sample demographics.

**Table 1.** Sample demographics. The table shows sample demographics for the MRI sample and the cognition sample. The total ethnicity numbers exceed total sample size as several subjects identified as belonging to more than one ethnicity group.

|  | **MRI sample** | **Cognition sample** |
| --- | --- | --- |
| **N** | 621 | 611 |
| **Age (years)** | 3.2-21.0 (mean= 12.4, SD= 4.8) | 3.2-21.0 (mean= 12.5, SD= 4.8) |
| **Sex** | Boys= 318, Girls = 303 | Boys=310, Girls = 301 |
| **Self-reported Ethnicity** | Hispanic/Latino = 140<br>Pacific Islander = 5<br>Asian = 88<br>African American = 102<br>Native American = 31<br>White = 426 | Hispanic/Latino = 136<br>Pacific Islander = 4<br>Asian = 86<br>African American = 100<br>Native American= 31<br>White = 420 |
| **Handedness** | Right handed= 532<br>Left handed= 60<br>Mixed =24<br>Not established = 3<br>Not available = 2 | Right handed = 523<br>Left handed= 60<br>Mixed = 24<br>Not established = 3<br>Not available = 1 |
| **Ever diagnosed with ADHD** | Yes = 48<br>No = 567<br>Not available = 6 | Yes = 48<br>No = 557<br>Not available = 6 |

### MRI acquisition and processing

Imaging data for our final sample was attained on 7 separate 3T scanners; 2 from GE medical systems (Signa HDx and Discovery MR750), and 5 from Siemens (TrioTim). Sex and age-ranges were largely overlapping across scanners (Figure 1, Supporting Table 1). To account for challenges associated with multisite imaging of children, an optimized imaging protocol was implemented across vendors and models to obtain similar contrast properties and image-derived quantitative measures (Jernigan et al., 2016). The protocol included a 3D T1w inversion prepared RF-spoiled gradient echo scan, and a 3D T2w variable flip angle fast spin echo scan (Jernigan et al., 2016), both acquired with prospective motion correction (PROMO) (White et al., 2010). Voxel sizes ranged from ≈ 0.9-1.2mm^3^. See Supporting Information for additional acquisition parameters, and care and safety procedures implemented for scanning children.

**Figure 1.**
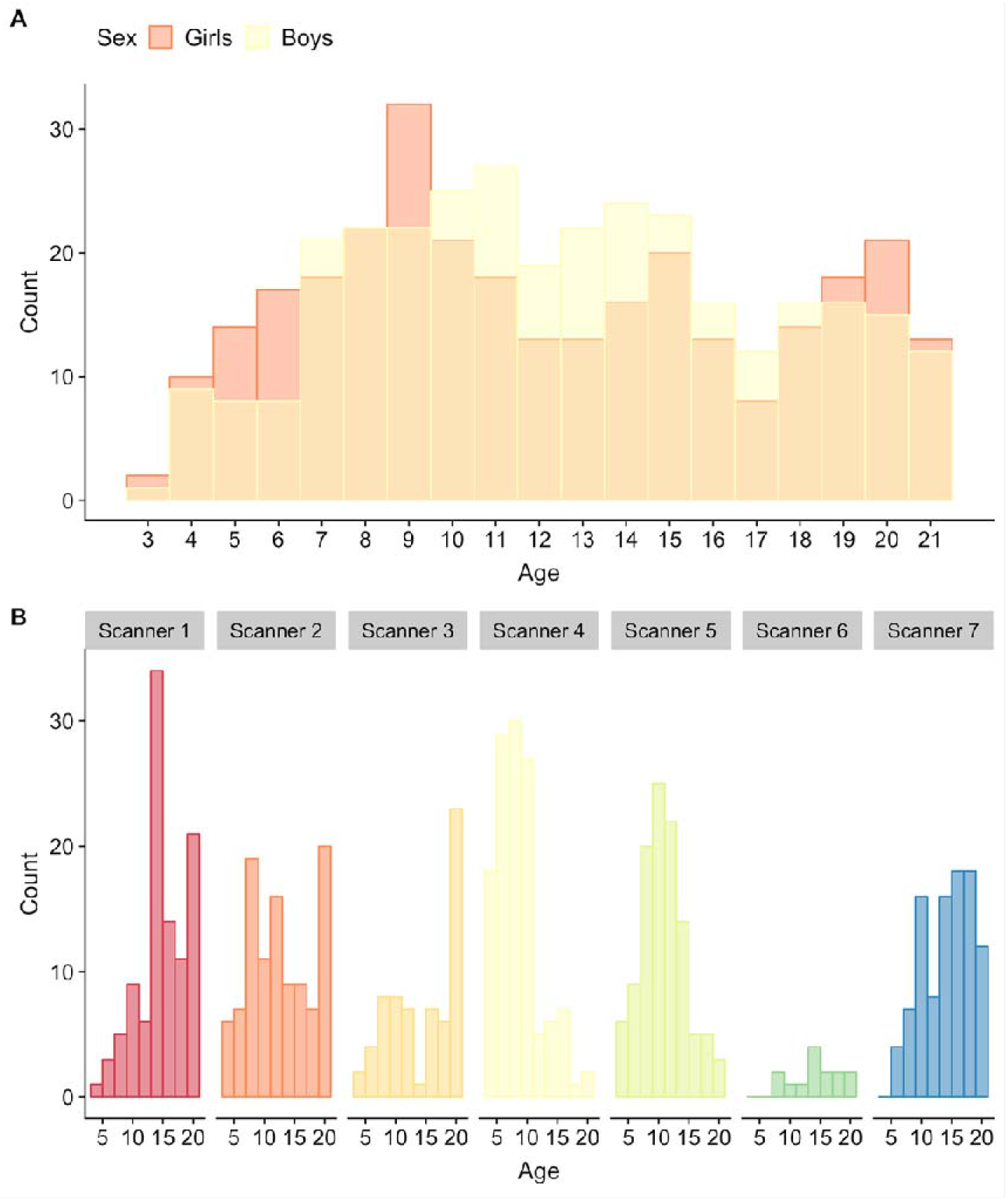
Age and sex distributions for the full MRI sample (n= 621). Plot A depicts age and sex distributions of the full MRI sample, while B depicts the age distribution for each scanner.

Imaging processing was performed using the three step Human Connectome Project (HCP) pipeline as described elsewhere (Matthew F. Glasser et al., 2013). In short, the “PreFreeSurfer” stage produces an undistorted native structural volume space, aligns the T1w and T2w image, performs bias field correction calculated from the product of both images, and registers the native- to the common coordinate MNI space using a rigid 6 degrees of freedom transform, derived from a 12 degrees of freedom affine registration. Based on FreeSurfer version 6, the “FreeSurfer” stage performs volume segmentation, and cortical surface reconstructions, including the “white”- (grey/white matter boundary), and “pial”- (grey/cerebrospinal fluid (CSF) boundary) surface, and registers the surfaces to a common template (fsaverage). Due to some cases exhibiting large cortical thickness and T1w/T2w ratio discrepancies between the left and right hemisphere following the default pipeline, we removed a flag (“t2pial”) within autorecon3, which when employed additionally uses the T2w sequence to reconstruct the pial surface. The “PostFreeSurfer” stage produces NIFTI volume- and GIFTI surface files, registers surfaces to the HCP standard mesh, and creates the final brain masks and T1w/T2w ratio maps (Matthew F. Glasser et al., 2013).

### T1w/T2w ratio maps

The creation of T1w/T2w ratio maps was based on methods described by M. F. Glasser and Van Essen (2011). A volume-to-surface mapping algorithm was applied to all voxel centers located within the cortical ribbon to achieve image division. For each vertex within the HCP pipeline surface “midthickness”, ribbon voxels were selected within a cylinder centered on the vertex, orthogonal to the surface, and with a height and radius equal to the local cortical thickness. To remove voxels with significant blood vessel- and CSF partial voluming effects, voxels were excluded if the T1w/T2w value exceeds ±1 SD of all values within the cortical ribbon. The remaining ribbon voxels were then averaged following a Gaussian weighted function to produce a single value for each vertex, while the medial wall was assigned values of zero (M. F. Glasser & Van Essen, 2011). The final output includes standard and bias corrected non-smoothed and smoothed (Full width at half maximum = 4mm) maps.

### Quality assessment of imaging data

We performed extensive quality assessment as multisite projects inevitably creates inconsistencies and since in-scanner head motion is more frequent in younger subjects. For subjects with both T1w and T2w data of sufficient resolution, all NIFTI images were processed through the automatic quality assessment pipeline MRIQC (Esteban et al., 2017). This program returns flagging of poor images as well as a numeric quality index based on measures of noise, spatial and tissue distribution, artifacts such as motion, and sharpness (Esteban et al., 2017). After visual inspection of flagged datasets by a single trained rater, subjects were either included, tagged for detailed re-assessment after T1w/T2w ratio map creation, or excluded due to poor image quality when there was no option to replace the image with another satisfactory run. The remaining subjects were subsequently processed through the HCP pipeline (Matthew F. Glasser et al., 2013). Finally, the same rater visually inspected all T1w/T2w ratio maps and excluded subjects with poor maps. See Supporting Information for in detail descriptions of quality assessments.

### Cognitive assessment

All subjects completed the computerized NIH Toolbox Cognition Battery (Akshoomoff et al., 2014; Weintraub, Bauer, et al., 2013; Weintraub, Dikmen, et al., 2013) which includes seven different tasks, designed to measure eight diverse cognitive abilities, and is appropriate for use in children as young as 3 years of age (Table 2, and Supporting Information). Test order was standardized across participants, and eight sum scores were calculated as described elsewhere (Akshoomoff et al., 2014). Due to missing cognitive data on specific tests, we imputed 78 sum score values in total, from 63 subjects in R, using the “mice” package (Buuren & Groothuis-Oudshoorn, 2011), with the default of 5 multiple imputations, and choosing the first imputation. As expected, there were high correlations between all sum scores and age (Pearson’s r ranging from 0.63 to 0.82), and we therefore residualized sum scores by linear and quadratic age by linear regression. Retrieving a single factor for general cognitive ability, conceptually similar to the IQ score, we then performed a principal component analysis (PCA) in R (https://www.r-project.org/) and the first factor, explaining 33.7% of the variance, was extracted and used as a measure of general cognitive ability. We changed the sign of the component so that higher factor loading would represent better performance. See Supporting Figure 1 and 2 for a bar plot of the percentage explained variance for each principal component, and the percentage contribution of each cognitive sum score to the first principal component, respectively. See Figure 2 for a correlation matrix between age, sex, general cognitive ability and individual cognitive sum scores.

**Table 2.** Overview of the NIH Toolbox Cognition Battery measures. The table shows cognitive tests in the order they were presented to subjects, the cognitive abilities assessed, and the abbreviations used in the current paper.

| **Test** | **Cognitive Ability** | **Abbreviation** |
| --- | --- | --- |
| Dimensional Change Card Sort Test | Cognitive flexibility | Cognitive flexibility |
| Flanker Inhibitory Control and Attention Test | Inhibitory control | Inhibition |
|  | Visual attention | Attention |
| Picture Sequence Memory Test | Episodic memory | Episodic memory |
| Pattern Comparison Processing Speed Test | Processing speed | Processing speed |
| Oral Reading Recognition Test | Oral reading skill | Reading |
| List Sorting Working Memory Test | Working memory | Working memory |
| Picture Vocabulary Test | Vocabulary knowledge | Vocabulary |

**Figure 2.**
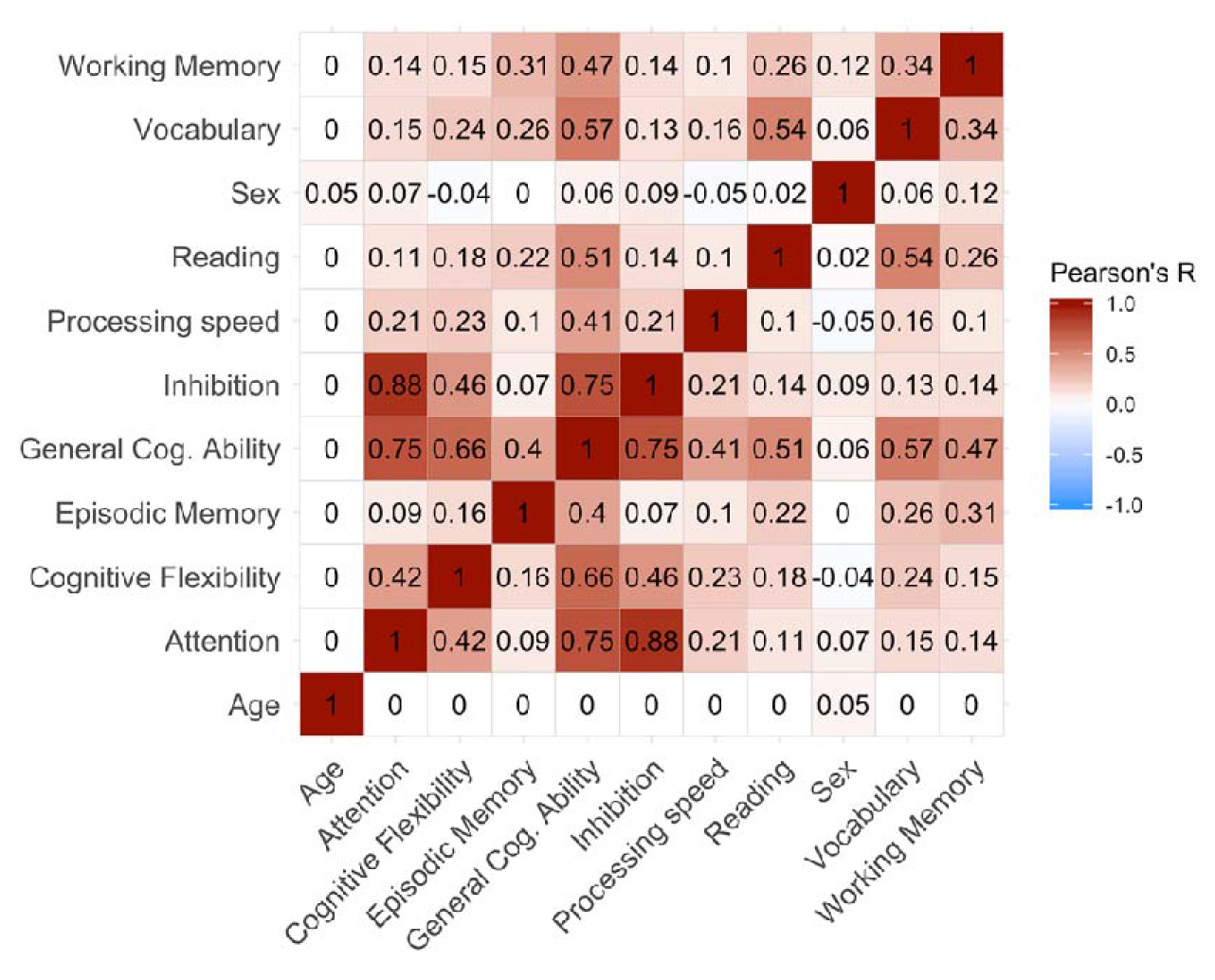
Correlation matrix for age, sex and cognition. The figure depicts Pearson’s correlation coefficients for age, sex, general cognitive ability (General Cog. Ability), and individual cognitive sum scores. The red spectrum represents positive correlations, while the blue spectrum represents negative correlations. Sum scores, and by extension general cognitive ability, are residualized by linear and quadratic age.

### Statistical analyses

We used general linear models (GLM), as implemented in the Permutation Analysis of Linear Models (PALM) toolbox (Winkler, Ridgway, Webster, Smith, & Nichols, 2014), to test for linear and quadratic (defined as polynomial of degree 2) effects of age on vertex-wise T1w/T2w ratio, covarying for sex and scanner. Second, we tested for main effects of sex on vertex-wise T1w/T2w ratio, covarying for linear and quadratic age and scanner, and then for interaction effects between sex and age, with main effect of sex as an additional covariate. Third, we performed vertex-wise analyses to assess the effects of general cognitive ability, as well as each of the eight specific sum scores, all in separate models on vertex-wise T1w/T2w ratio, each time with sex, linear and quadratic age, sex*age and scanner as co-variates. Finally, we examined interaction effects between age and general cognitive ability on T1w/T2w ratio, with sex, linear and quadratic age, sex*age, general cognitive ability and scanner as co-variates. To assess statistical significance we used 10,000 permutations and family wise error (FWE) correction with threshold-free cluster enhancement (TFCE) (Smith & Nichols, 2009) corrected across each contrast (as PALM performs one tailed analyses) and a significance threshold of p<0.05, after correcting for both hemispheres.

To regionally summarize relevant vertex-wise results, we averaged our t-statistic surface maps within 34 regions of interest (ROIs) for both the left and right hemisphere. ROIs were defined by Freesurfer’s fsaverage Desikan-Killiany Atlas parcellation (Desikan et al., 2006; Fischl et al., 2004) which was converted to GIFTI format and mapped to the HCP common mesh. To further visualize relevant PALM results, we used individual parcellations and locally weighted scatterplot smoothing (loess) and ggplot2 (Wickham, 2009) in R (https://www.r-project.org/).

## Results

### T1w/T2w mean ratio map

Figure 3 shows the mean T1w/T2w ratio map across all subjects. Briefly, the anatomical signal distribution corresponded well with previous T1w/T2w maps (M. F. Glasser & Van Essen, 2011), showing a specific regional pattern in which sensorimotor regions, such as pre- and post-central gyri and visual cortex, showed relatively higher T1w/T2w ratio, while the frontal lobe and temporal pole showed relatively lower T1w/T2w ratio.

**Figure 3.**
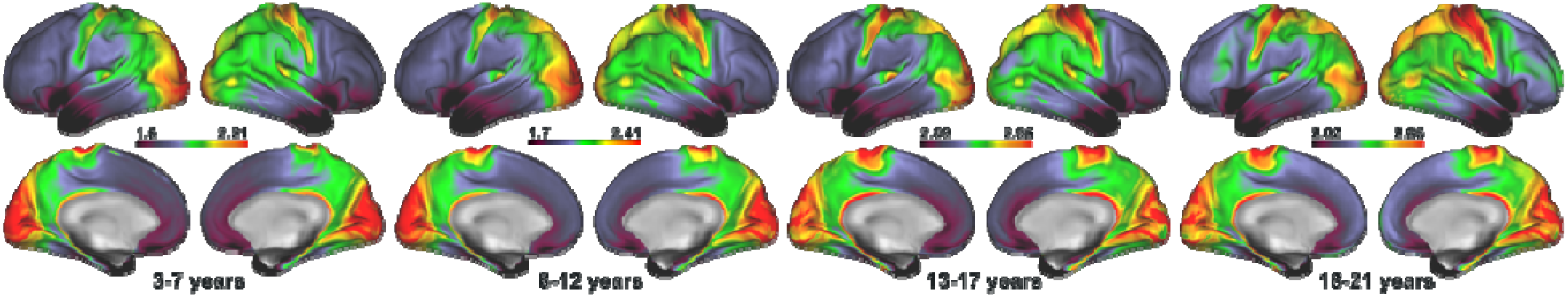
Age binned mean T1w/T2w ratio maps. The figure depicts mean T1w/T2w ratio maps from the full MRI sample (n= 621), devided into 4 age bins of 3-7 years (n= 120), 8-12 years (n= 219), 13-17 years (n= 168), and 18-21 years (n= 114). Warm colors represent regions with higher T1w/T2w ratio, while cold colors represent regions with lower T1w/T2w ratio.

For regional box plots of T1w/T2w ratio intensity spread across scanners see Supporting Figure 3. In short, all but one scanner showed similar T1w/T2w ratio intensity spreads.

### Associations between T1w/T2w ratio and age

Permutation testing revealed a near global positive association between linear age and T1w/T2w ratio, indicating higher ratio in widespread regions with higher age when controlling for sex (Figure 4). There was a general posterior to anterior increasing gradient in association strength, with highest mean t-statistics found in precentral, caudal middle frontal, pars triangularis, and postcentral regions within the left hemisphere, and rostal middle frontal, caudal middle frontal, precentral, and paracentral regions within the right hemisphere (Table 3). Sub-regions within the medial occipital lobe and minor medial parietal and frontal regions did not show significant age associations.

**Figure 4.**
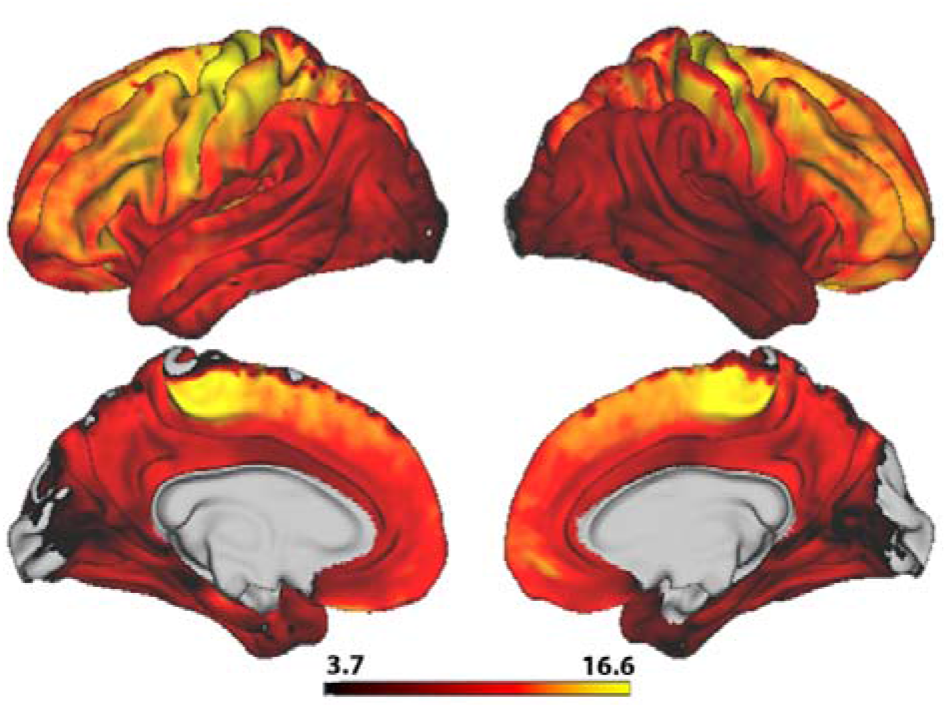
Vertex-wise associations between T1w/T2w ratio and age. The figure depicts a t-statistics map, masked by familywise error corrected p values across contrasts and thresholded at a minimum −log_p_ of 1.6 to correct for both hemispheres. Warm colors represent a postive age association.

**Table 3.**
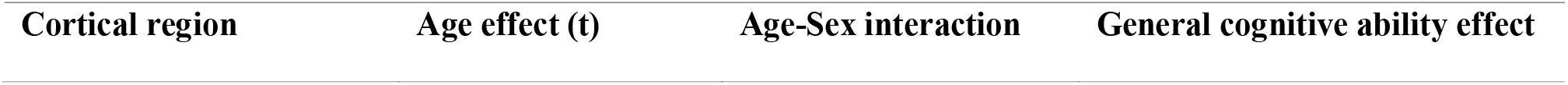
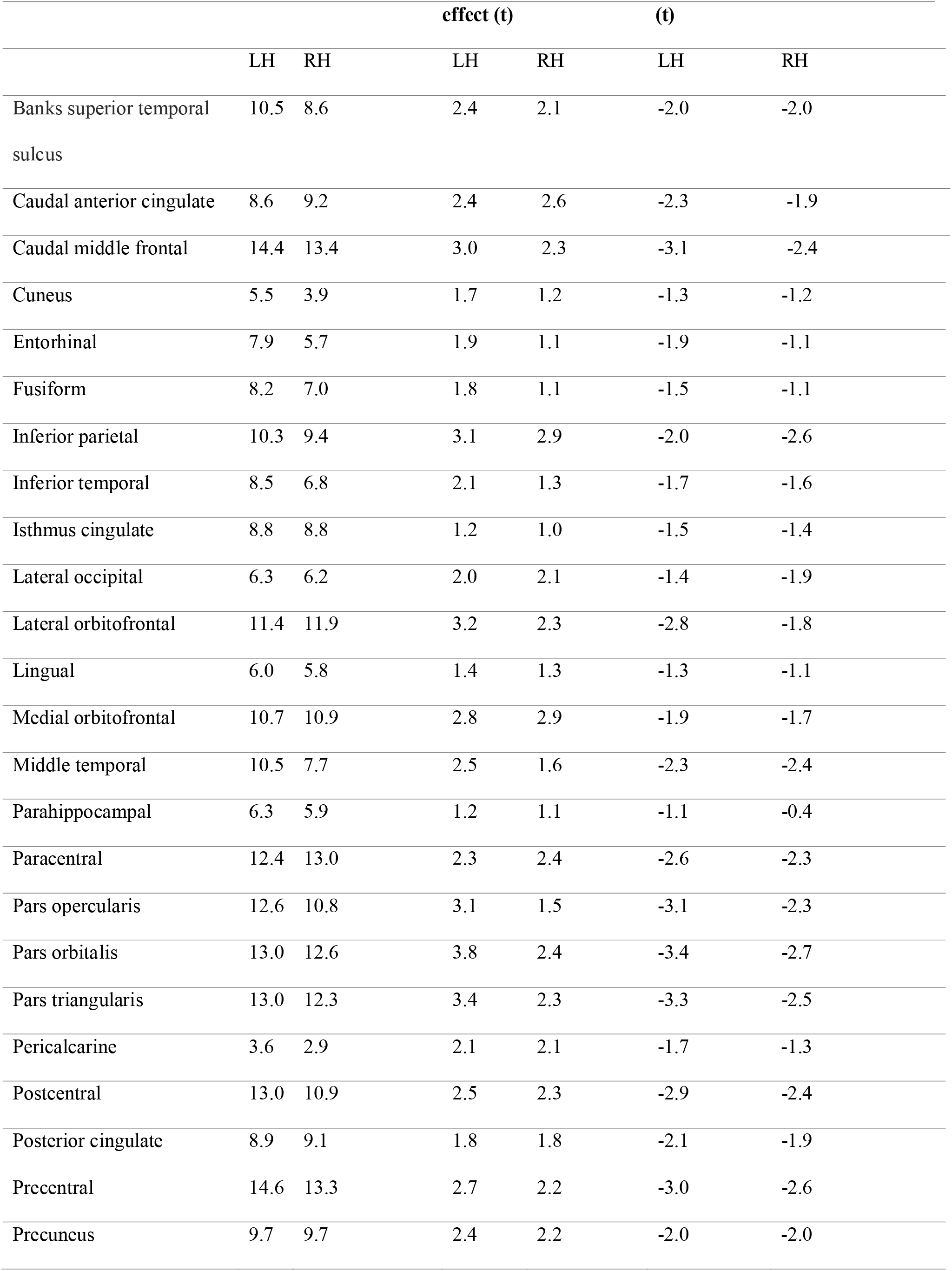
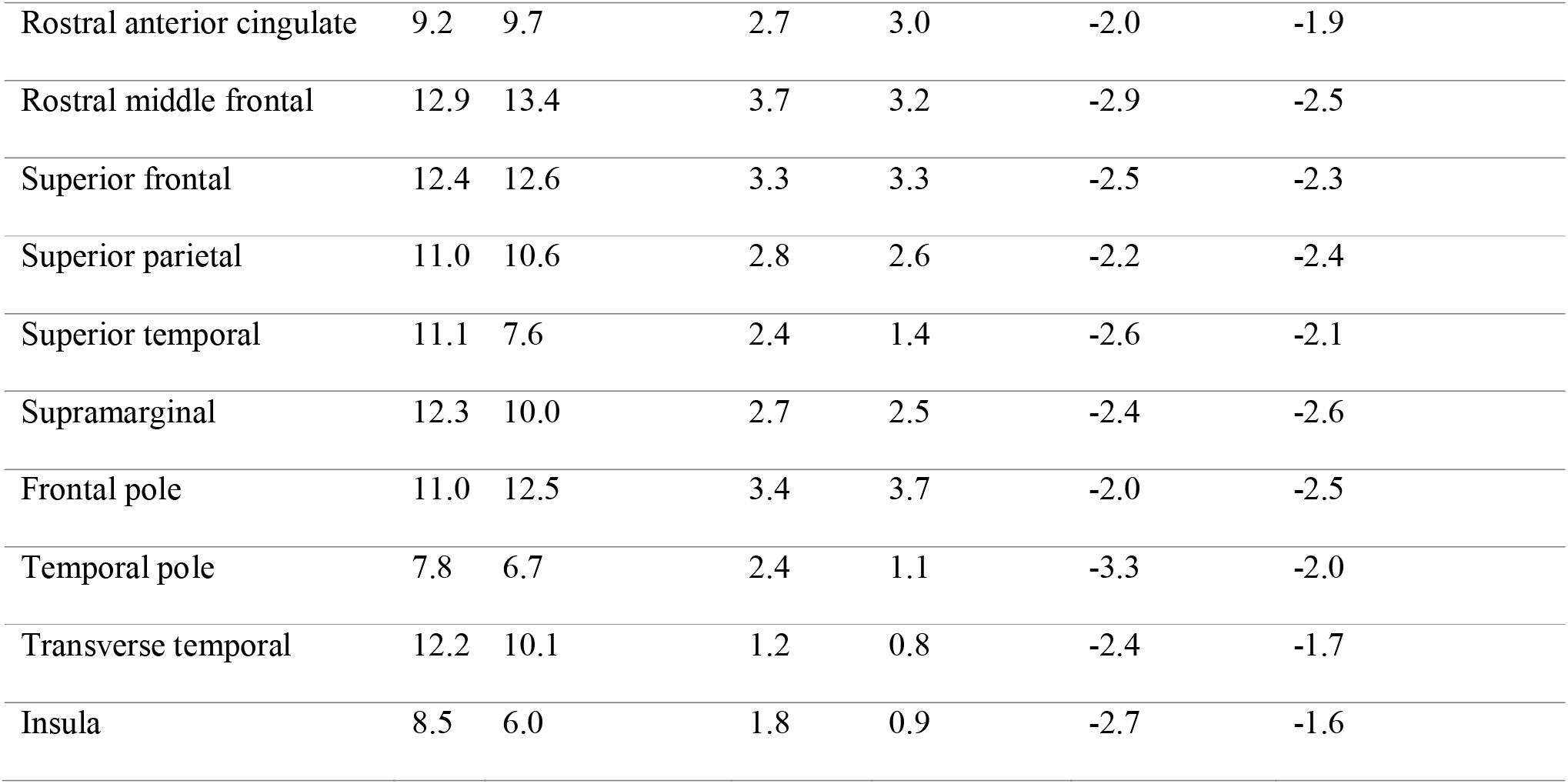
ROI based mean t-statistics of T1w/T2w ratio associations. The table shows each of the 34 ROIs, and the mean t-statistics within each region for the association between T1w/T2w ratio and age, age*sex and general cognitive ability within left and right hemisphere separately.

There were no statistically significant associations between T1w/T2w ratio and quadratic age, controlling for linear age and sex. Corresponding with our vertex-wise results, ROI-based plots with non-parametric loess curves showed steady increases of T1w/T2w ratio with higher age (Figure 5).

**Figure 5.**
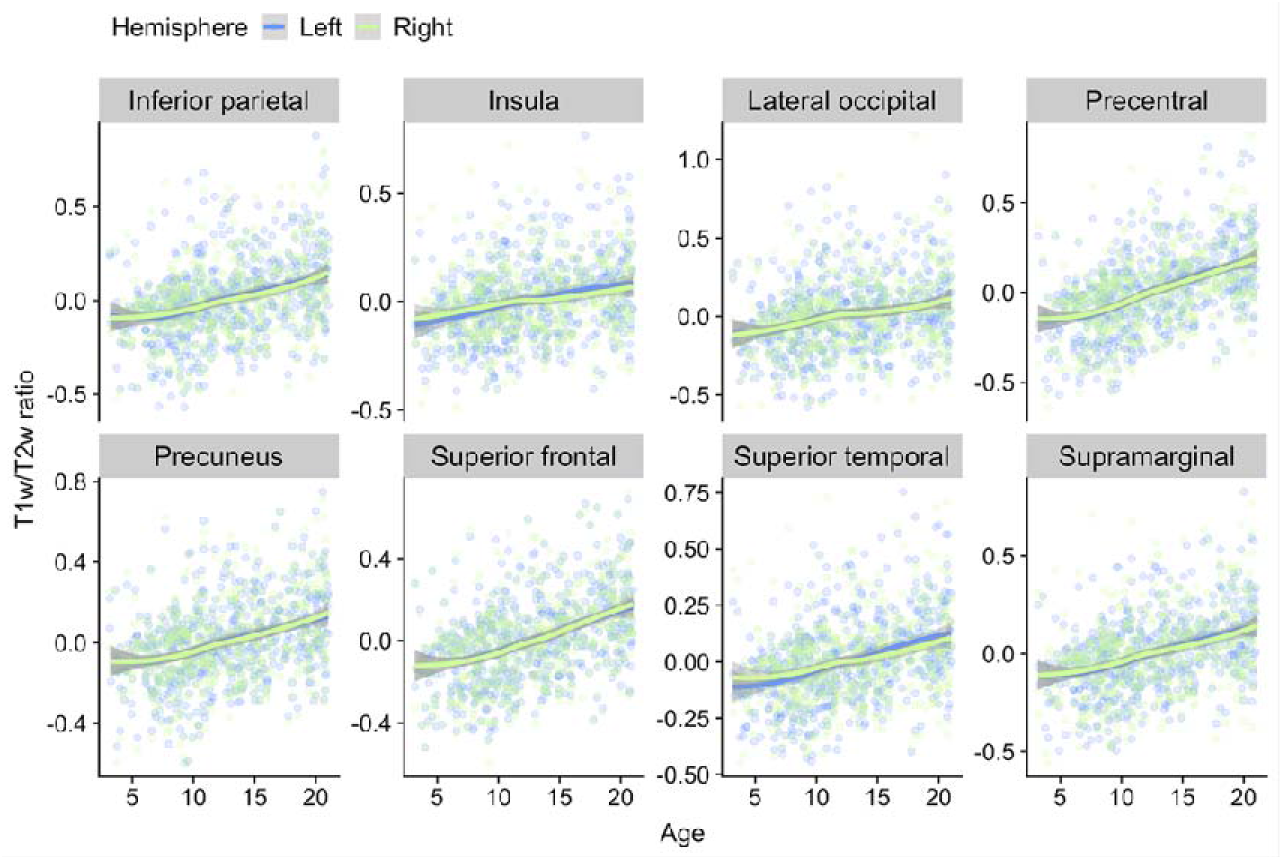
Regional loess visualizations of associations between T1w/T2w ratio and age. The plots show age plotted against mean T1w/T2w ratio within a selection of regions from different cortical lobes. Data points and smooth splines are color coded by hemisphere. T1w/T2w ratio values are residualized by sex and scanner.

See Supporting Figure 4 for correlations between regional T1w/T2w ratio and age across scanners. In short, scanners showed relatively similar correlation coefficients.

### Associations between T1w/T2w ratio and sex

Permutation testing revealed no statistically significant main effect of sex on vertex-wise T1w/T2w ratio. There were, however, significant positive interaction effects between sex and age (Figure 6). Effects were more widespread in the left compared to the right hemisphere with highest mean t-statistics found in pars orbitalis, rostral middle frontal, pars triangularis and frontal pole in the left hemisphere, and frontal pole, superior frontal, rostral middle frontal and rostral anterior cingulate in the right hemisphere (Table 3). Corresponding with our vertex-wise results, frontal ROI-based plots showed a pattern where boys within the age-range of approximately 15-21 years had relatively higher T1w/T2w ratio than girls (Figure 7).

**Figure 6.**
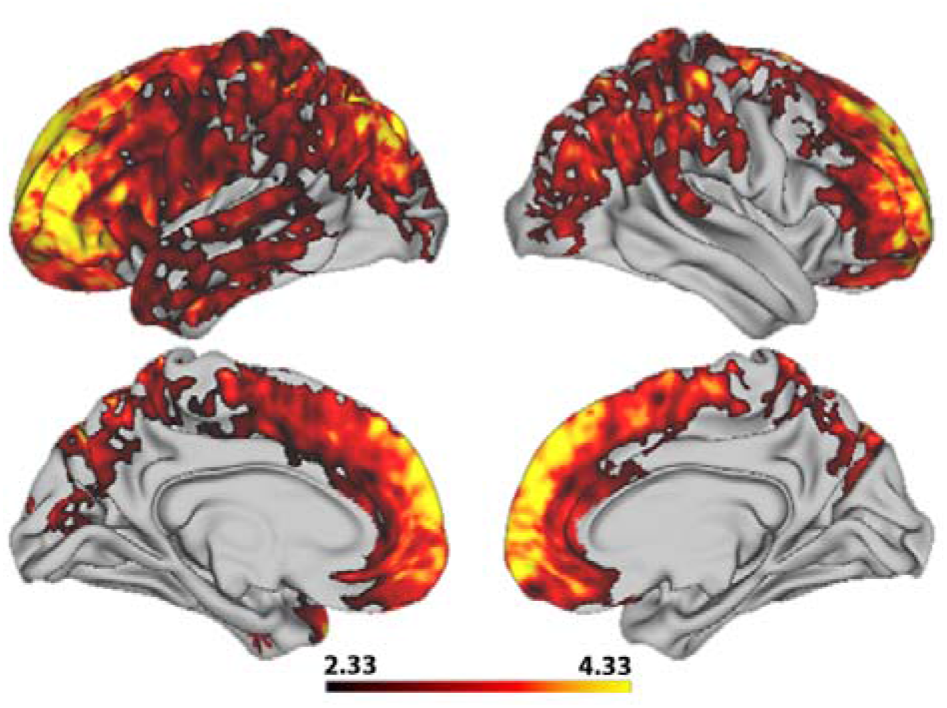
Vertex-wise associations between T1w/T2w ratio and the interaction between age and sex. The figure depicts a t-statistics map, masked by familywise error corrected p values across contrasts and thresholded at a minimum −log_p_ of 1.6 to correct for both hemispheres. Warm colors represent a postive interaction effect.

**Figure 7.**
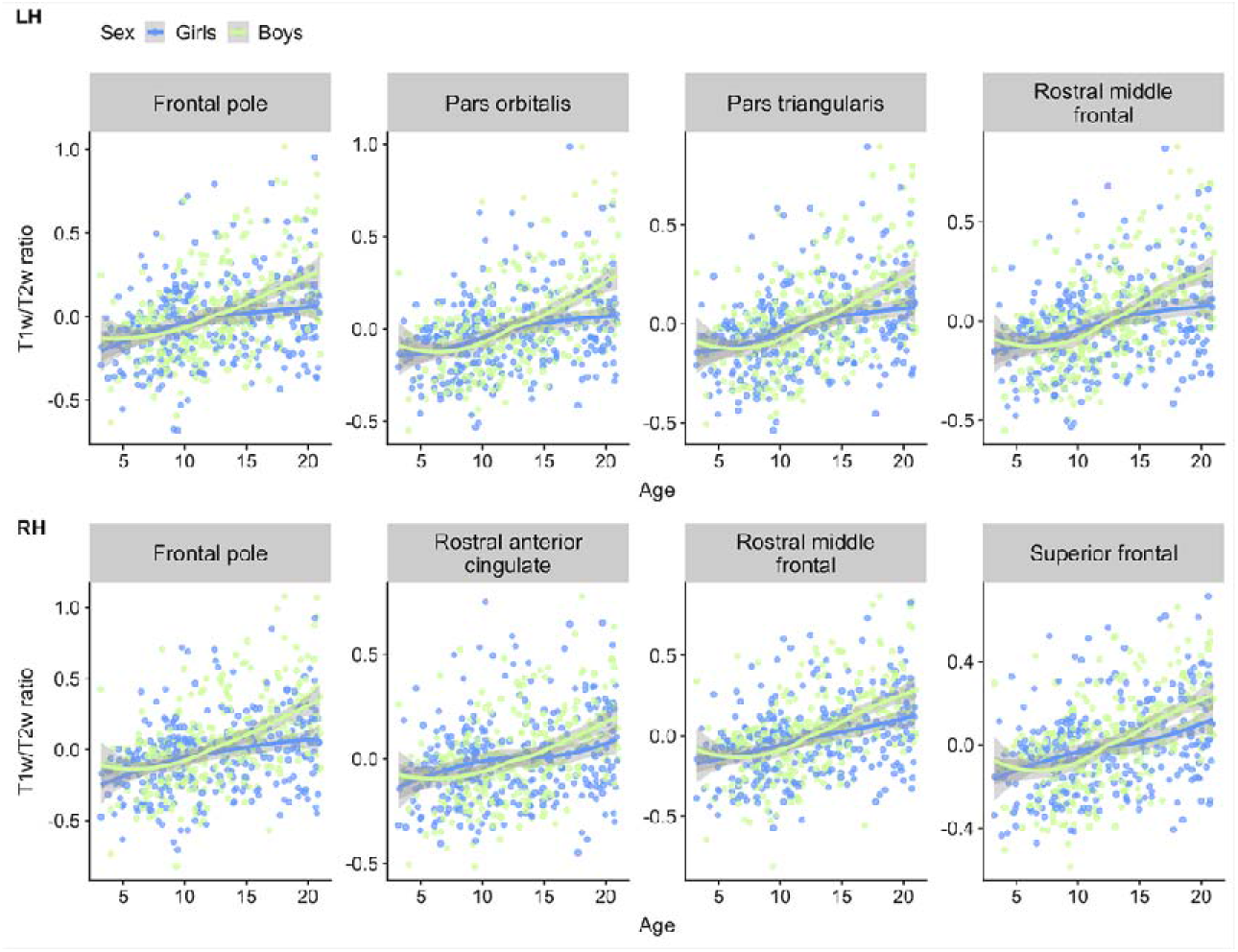
Visualizations of T1w/T2w ratio versus age color coded for sex. The plots shows age plotted against mean T1w/T2w ratio within frontal regions, selected based on the highest regional age*sex mean t-statistics, with data points and smooth splines color coded by sex. T1w/T2w ratio values are residualized by scanner.

### Associations between T1w/T2w ratio and cognition

Permutation testing revealed a negative relationship between T1w/T2w ratio and general cognitive ability in the frontal lobe extending into left temporal pole and right parietal regions, with more widespread associations in left compared to the right hemisphere (Figure 8). This indicates that lower T1w/T2w ratio was associated with better cognitive performance. Strongest negative associations were found in pars orbitalis, pars triangularis, temporal pole and caudal middle frontal regions in the left hemisphere, and pars orbitalis, inferior parietal, precentral and supramarginal regions in the right hemisphere (Table 3).

**Figure 8.**
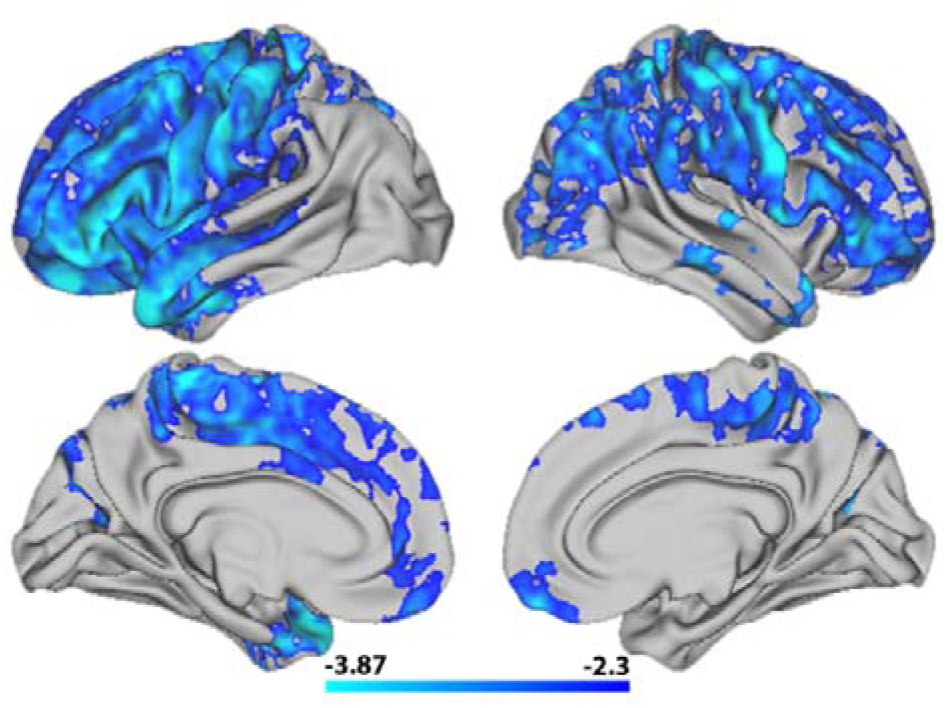
Vertex-wise associations between T1w/T2w ratio and general cogntive ability. The figure depicts a t-statistics map, masked by familywise error corrected p values across contrasts and thresholded at a minimum −log_p_ of 1.6 to correct for both hemispheres. Cold colors represent a negative association between T1w/T2w ratio and general cognitive ability.

Permutation testing revealed no significant interaction effects between general cognitive ability and age on T1w/T2w ratio. Corresponding with our vertex-wise results, ROI-based plots indicated that higher general cognitive ability was associated with regionally lower T1w/T2w ratio across the age-range (Figure 9). Moreover, 5 specific cognitive sum scores yielded regional negative associations with T1wT2w ratio, with varying cluster extension after correction across hemispheres, largely overlapping with the regions reported for general cognitive ability (Figure 10). Inhibition and attention showed small clusters within the left hemisphere, with inhibition covering precentral and postcentral sub regions, and attention covering the postcentral region only. While vocabulary only showed associations in small frontal clusters within the left hemisphere, the vertex-wise pattern for reading was widespread and highly similar to the pattern found for general cognitive ability, with frontal associations stronger in the left as compared to the right hemisphere. Finally, working memory showed regional associations within right hemisphere only, mainly in parietal regions, covering highly similar areas as general cognitive ability. There were no significant associations between T1w/T2w ratio and cognitive flexibility, episodic memory and processing speed.

**Figure 9.**
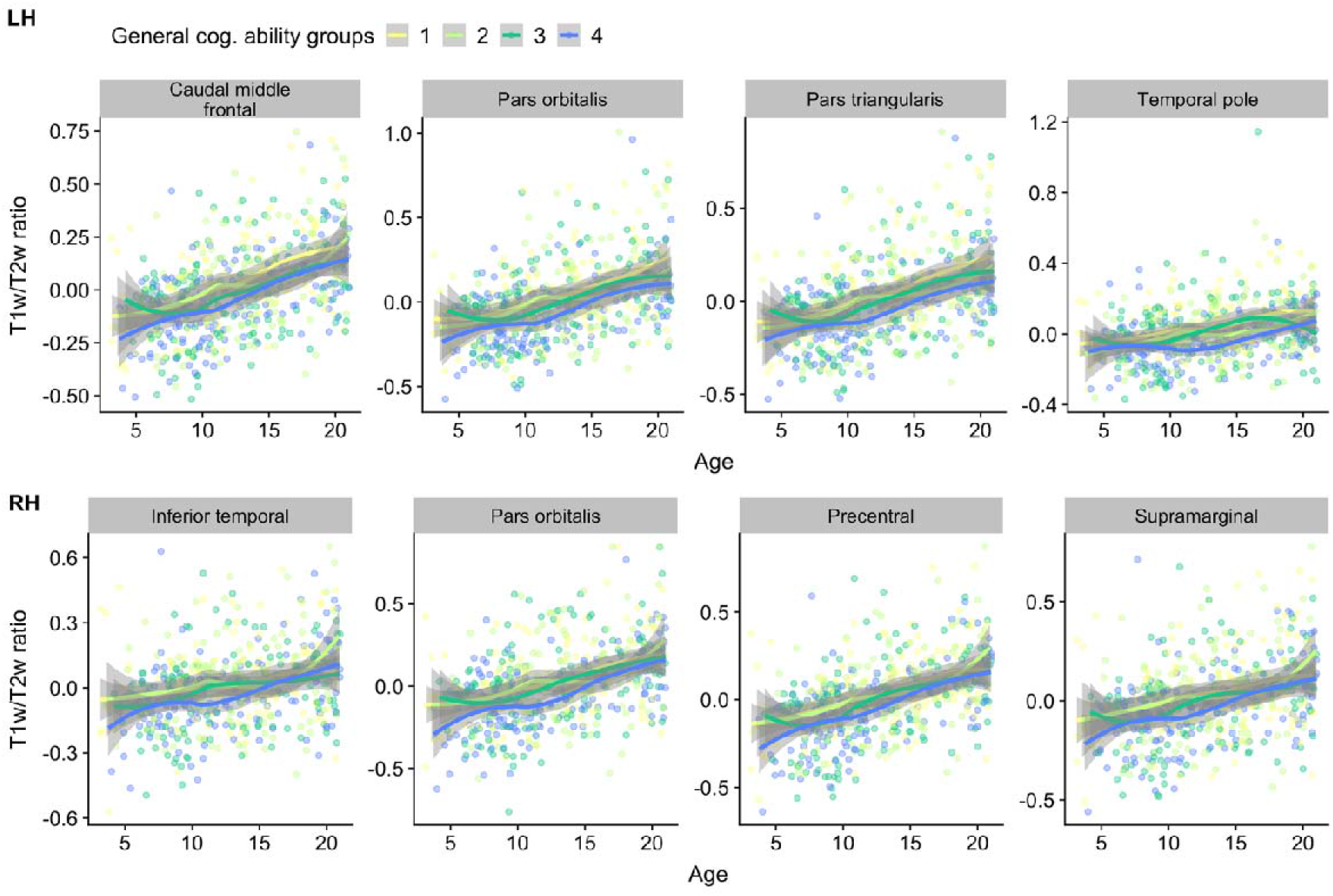
Visualizations of T1w/T2w ratio versus age, color coded by general cognitive ability. The plots depict age plotted against mean T1w/T2w ratio within frontal regions, selected based on the the highest regional mean t-statistics, with data points and smooth splines color coded by general cogntive ability, resulting in 4 equal qunatiles with worst (yellow) to best (blue) cognitive performers. T1w/T2w ratio is sex and scanner residualized, while general cognitive performance is residualized by linear and quadratic age.

**Figure 10.**
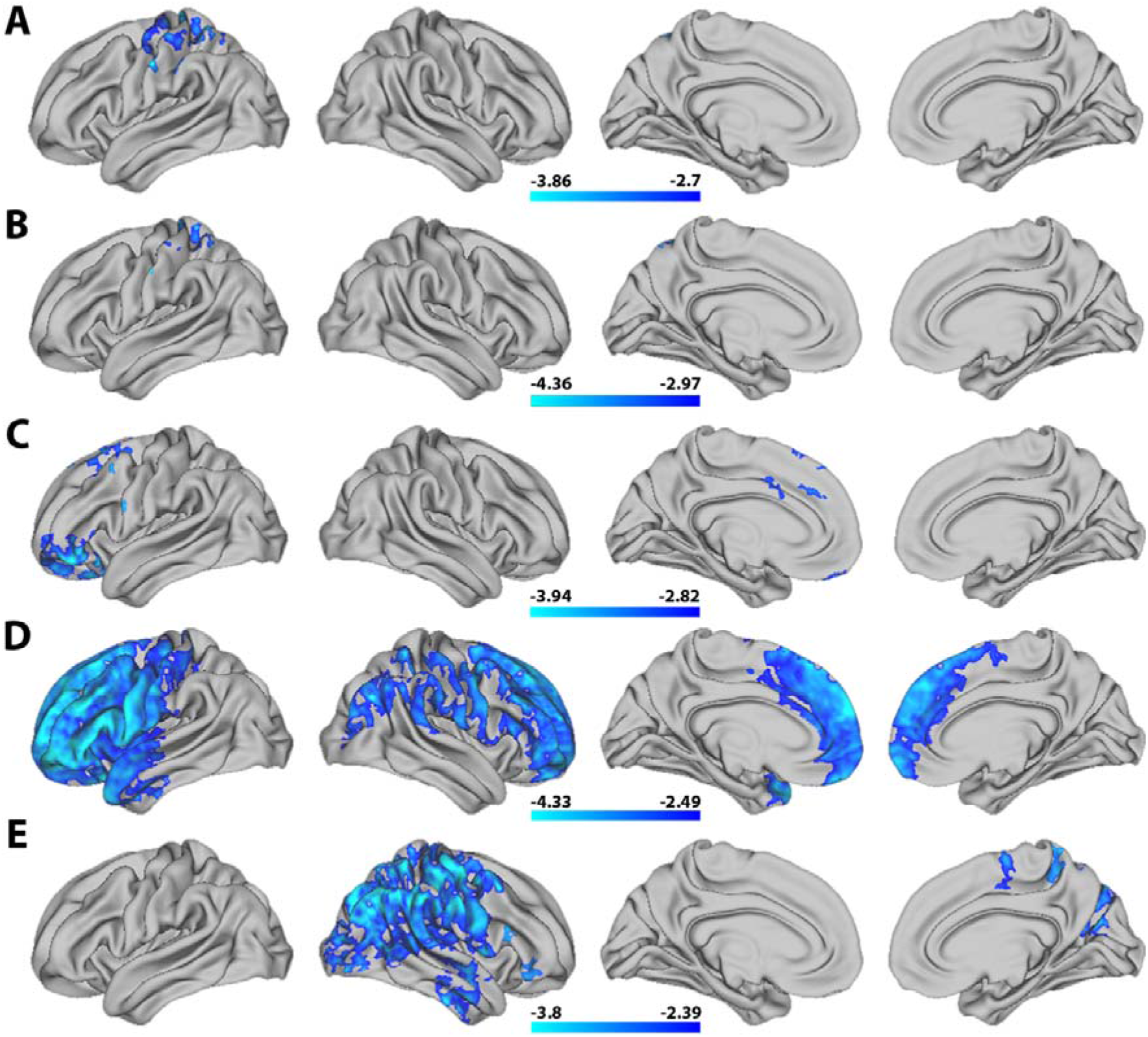
Vertex-wise associations between T1w/T2w ratio and individual cognitive sum scores. The figure depicts t-statistics maps from top to bottom for (A) inhibition, (B) attention, (C) vocabulary, (D) reading, and (E) working memory. The maps are masked by familywise error corrected p values across contrasts and thresholded at a minimum −log_p_ of 1.6 to correct for both hemispheres. Cold colors represent a negative association between T1w/T2w ratio and each sum score.

See Supporting Figure 5 for ROI based T1w/T2w ratio spread for individuals with left and right hand dominance. In short, spreads were highly similar across hemispheres.

## Discussion

The cerebral cortex is the center for most of our evolutionarily unique cognitive abilities (Geschwind & Rakic, 2013), and its maturation is therefore an essential part of human brain development, underpinning major environmental adaptations and vast cognitive improvements (Casey et al., 2005). In a large sample of youths aged 3-21 years we found a global age-related increase of T1w/T2w ratio across the brain surface, with some regional and sex related variation. Intriguingly, across individuals, T1w/T2w ratio was inversely associated with general and several specific cognitive abilities, mainly in anterior cortical regions.

The first goal of the present study was to investigate associations between T1w/T2w ratio and age. As hypothesized, we found a near global positive association, with a continuous age-related increase of T1w/T2w ratio from early childhood and into young adulthood. Our results converge with the few prior studies on T1w/T2w ratio in children, adolescents and young adults (Håkon Grydeland et al., 2013; Shafee et al., 2015). Grydeland et al. (2013) reported that for their youngest age group, consisting of 8 to 19 year old subjects, there was a positive linear association between T1w/T2w ratio and age across large portions of the cerebral cortex, with strongest association in posterior frontal, parietal, and temporal cortices. We too found our strongest associations in posterior frontal regions extending into parietal regions. In contrast to the present results, Grydeland et al. did not find age-associations within parts of anterior frontal and insular cortices, but this discrepancy could be due to greater sample size and the inclusion of younger children in the current study.

Our analyses revealed no significant main effects of sex on T1w/T2w ratio, but a positive age by sex interaction indicated that boys in late adolescence had relatively higher T1w/T2w ratio than girls in the frontal lobe and parts of parietal lobe. No previous studies have to our knowledge specifically investigated the relationship between T1w/T2w ratio and sex, but several studies have tested for sex differences in cortical morphometry and morphometric development. Although there are inconsistencies in this literature, recent longitudinal results indicate larger cortical volumes and areas in boys, but few differences in cortical thickness, and limited sex differences in the estimated rates of cortical morphometric development (Vijayakumar et al., 2016; Wierenga et al., 2019). A recent study investigating a longitudinal sample of 14 to 24 year old youths, using magnetization transfer (MT), an imaging measure linked to myelination, reported main effects of sex, with females having regionally higher MT than males (Ziegler et al., 2019). Less optimal cross sectional analyses of the same data, yielded a similar sex-age interactional pattern to the one reported in the current study, within right angular gyrus. Therefore, future longitudinal studies on T1w/T2w ratio are needed for more direct investigations. Moreover, as several studies have reported sex differences in morphometric brain variation beyond limited mean effects (Wierenga et al., 2019; Wierenga et al., 2018) future studies could investigate whether boys show higher variability in T1w/T2w ratio than girls.

Few previous studies have explored the behavioral and cognitive relevance of T1w/T2w ratio during development. We found a negative association between T1w/T2w ratio and general cognitive ability, mainly in anterior regions. We also found corresponding associations regarding both direction and cumulative regional extension between T1w/T2w ratio and the specific cognitive abilities: inhibition, attention, reading, vocabulary and working memory. Comprehensive interpretations of structural brain-behavior relations should be done cautiously, as such relations are likely complex and have shown poor replication rates (Boekel et al., 2015; Kharabian Masouleh, Eickhoff, Hoffstaedter, & Genon, 2019). Nevertheless, reading and vocabulary surface maps showed left hemispheric dominance, in concordance with the general understanding that language processing is mainly located in the left hemisphere for right-handed individuals (Saur et al., 2008). The working memory surface map is also partly in line with previous reports of a central role for a front-parietal network (Salazar, Dotson, Bressler, & Gray, 2012; Tamnes et al., 2013). We did not find an interaction between cognitive ability and age on T1w/T2w ratio.

Intriguingly, this indicates that beyond developmental improvements, children and adolescents with high general- and specific cognitive performance have regionally lower T1w/T2w ratio especially within the frontal lobe. Considering the positive T1w/T2w ratio - age association, this result is surprising. However, it does converge with several studies reporting negative or otherwise unexpected direction of associations between diffusion metrics and cognitive skills in children (Dougherty et al., 2007; Huber, Henriques, Owen, Rokem, & Yeatman, 2019; Travis, Adams, Kovachy, Ben-Shachar, & Feldman, 2017). Our results also mostly concord with the very limited prior literature on T1w/T2w ratio and cognition in youth. Grydeland et al. (2013) investigated the relationship between T1w/T2w ratio and performance variability in response time in youths and adults. Although the general conclusion, based on the combined lifespan sample, was that higher T1w/T2w ratio was associated with greater performance stability, the opposite relationship was reported for the young subgroup of 8 to 19 year old subjects in clusters within occipital and parietal regions. However, a recent study by Vandewouw et al. (2019) found lower T1w/T2w ratio within occipital, parietal and subcortical regions in a small group of four year old preterm children as compared to children born full term. Across the two groups, they also found positive associations between T1w/T2w ratio in parietal lobes and intelligence, and expressive and receptive language ability.

The biological underpinnings of the T1w/T2w ratio is debated and likely complex. On the one hand, intracortical myelin content inversely co-varies with the T1w and T2w signal intensity (M. F. Glasser & Van Essen, 2011), through differences in lipids (Koenig, 1991), water concentration (Miot-Noirault, Barantin, Akoka, & Le Pape, 1997), and iron which in the cortex is strongly co-located with myelin (Fukunaga et al., 2010). The T1w- to T2w division also mathematically cancels most scanner related intensity biases, while increasing contrast (M. F. Glasser & Van Essen, 2011), and T1w/T2w ratio concurs with post mortem data (Nakamura et al., 2017). On the other hand, recent attempts have not revealed close links between T1w/T2w ratio and either myelin-related genes or more established myelin measures (Hagiwara et al., 2018; Ritchie et al., 2018; Uddin et al., 2019), demonstrating that it is not a straightforward myelin proxy. Nevertheless, our T1w/T2w ratio - age results are in line with the known protracted maturational timing and directional pattern of intracortical myelination (Miller et al., 2012; Yakovlev & Lecours, 1967). The observed negative associations between T1w/T2w ratio and cognition are less intuitive if interpreted through a cortical myelin perspective. One could speculate that having excess levels of cortical myelin beyond a certain normative developmental range, is disadvantageous as myelin contains factors associated with inhibition of neurite growth. Excess myelin might consequently foster a permanent inhibitory environment for synapse formation and lower neuronal plasticity (Snaidero & Simons, 2017). Another possibility, as suggested by several diffusion studies (Dougherty et al., 2007; Huber et al., 2019; Travis et al., 2017) is that, in children, other tissue factors besides myelin content may account for variations in cognitive skills. As the frontal lobe contains low myelin content and shows protracted maturation, there might be distinct tissue properties underlying our cognitive- and age related findings. On a related note, recent findings suggest that MRI based apparent cortical thinning in youth is partly due to an increasing underestimation of the true cortical thickness. The rationale is that axonal myelination occurring predominantly within deep cortical layers brightens its appearance resulting in a misclassification of these voxels as white matter, consequently shifting the grey/white boundary outward (Natu et al., 2019; Sowell et al., 2004; Walhovd, Fjell, Giedd, Dale, & Brown, 2017; Westlye et al., 2009; Whitaker et al., 2016). T1w/T2w ratio is calculated within the cortical ribbon, a reconstruction in part based on the grey/white boundary. One could speculate that developmental processes in high cognitive functioning youths increase T1w intensities within deep cortical layers, thereby shifting such voxels from the cortical ribbon and into white matter, consequently giving rise to a thinner but also “darker” cortical ribbon as compared to their poorer cognitive performing peers. If so, beyond typical age related changes (which indeed could in reality also be larger) lower T1w/T2w ratio would be associated with higher cognitive functioning in youth.

We did not find associations between T1w/T2w ratio and age in sub regions of medial occipital lobe and minor medial parietal and frontal regions, probably due to removal of the t2pial flag during the HCP pipeline (see Supporting Information). Other study limitations include the cross-sectional design, which is a sub-optimal and indirect way to investigate maturation and development. Moreover, imaging data was attained on several different scanners. Although multisite initiatives to pool data is essential to battle challenges of underpowered neuroimaging studies, these strategies come with new challenges and limitations, including scanner and sequence related variability. In the current study we chose to statistically control for scanner in all analyses and age and sex were similar across scanners. Scanner related effects will still most likely confound analyses, including possible non-linear effects. Nevertheless, supplementary analyses indicated similar T1w/T2w ratios across all but one scanner, and similar correlations between regional T1w/T2w ratios and age. Finally, during development T1w/T2w ratio might be affected by an outward-shifting of the grey/white boarder, and future studies are needed to directly investigate the relations between cortical morphometric and microstructural measures.

To conclude, our results indicated that cortical T1w/T2w ratio almost globally increases from early childhood until young adulthood, corresponding with established maturational patterns of intracortical myelination. We also found that T1w/T2w ratio was inversely associated with measures of both general and several specific cognitive abilities, especially in anterior cortical regions. Future studies, preferably longitudinal, are needed to validate the robustness of these developmental patterns and links with individual cognitive differences and developmental psychopathology.

## Acknowledgments and disclosures

This work was supported by the Department of Psychology, University of Oslo (LBN and CKT), the Research Council of Norway (223273, 249795, 230345), the South-Eastern Norway Regional Health Authority (2019069, 2014097, 2016083), and the European Commission’s 7th Framework Programme (602450, IMAGEMEND). Data collection and sharing was funded by the Pediatric Imaging, Neurocognition and Genetics Study (PING) (National Institutes of Health Grant RC2DA029475). PING is funded by the National Institute on Drug Abuse and the Eunice Kennedy Shriver National Institute of Child Health & Human Development. PING data are disseminated by the PING Coordinating Center at the Center for Human Development, University of California, San Diego. Data were also in part provided by the Human Connectome Project, WU-Minn Consortium (Principal Investigators: David Van Essen and Kamil Ugurbil; 1U54MH091657) funded by the 16 NIH Institutes and Centers that support the NIH Blueprint for Neuroscience Research; and by the McDonnell Center for Systems Neuroscience at Washington University. A preprint is published on bioRxiv.org. The authors report no biomedical financial interests or potential conflicts of interest.

