## Supplemental materials for "Maturation of cortical microstructure and cognitive development in childhood and adolescence: a T1w/T2w ratio MRI study"

**Supporting Materials and Methods**

### Care and safety procedures during image acquisition

The current sample included very young children, and it should therefore be explicitly stated that no subjects were sedated for imaging (Brown et al., 2012). Before scanning, subjects were accompaniment by a parent or a technician into the scanning environment and exposed and habituated to the MRI machine before image acquisition. It was also possible to take breaks in between scans, or receive other behavioral support if needed. During scanning, all subjects were asked to lay as still as possible and the youngest subjects were given additional head padding. During the structural protocol, participants were presented with a movie of their choice with sound provided via headphones (Brown et al., 2012).

*MRI acquisition*

On each scanner, standardized multiple modality high-resolution protocols identical or nearly identical to the pulse sequence parameters implemented at UC San Diego were installed. At UCSD, data were obtained on a GE 3T Signa HDx scanner and a 3T Discovery 750x scanner (GE Healthcare, Waukesha, WI) using eight-channel phased array head coils. The protocol included a sagittal 3D inversion recovery spoiled gradient echo (IR-SPGR) T1-weighted volume which was optimized for maximum gray/white matter contrast (TE = 3.5 ms, TR = 8.1 ms, TI = 640 ms, flip angle = 8°, receiver bandwidth = ± 31.25 kHz, FOV = 24 cm, freq = 256, phase = 192, slice thickness = 1.2 mm). It also included a sagittal 3D cube T2-weighted volume (TE = 69.3 ms, TR = 1500 ms, echo train = 40, FOV = 24 cm, freq = 256, phase = 192, slice thickness = 1.2 mm). The scanning durations was 8:05 and 4:25 minutes for the T1w and T2w scan respectively (Brown et al., 2012). Further details are described elsewhere (Brown et al., 2012; Jernigan et al., 2016; White et al., 2010).

*MRI quality assessment*

From 998 available subjects with available MRI and demographic data, several were first directly excluded due to lack of a T2 sequence, for having unsatisfactory T1 or T2 resolution > 1.2mm voxel size in any direction, and for having multiple runs within the same folder, highly suspected of being different individuals (n=322). All NIFTI images including several duplicate images, unique re-runs from the same session, and images from the same subject but at different sessions i.e. after re-positioning in the scanner, were then processed through the quality assessment pipeline MRIQC (Esteban et al., 2017). After visual inspection of flagged images (n=99), by a single experienced rater, flagged subjects were either included, tagged for re-assessment after T1w/T2w ratio map creation, or excluded due to poor image quality (n=17) when there was no option to replace the image with another satisfactory run. For non-flagged subjects with several images, the highest quality sequence was chosen based on having superior “quality index” number which is outputted from MRIQC.	The remaining subjects were subsequently processed through the full Human Connectome Project (HCP) pipeline (Glasser et al., 2013) of which a few (n= 4) subjects failed completion and were excluded. Quality assessment of the T1w/T2w ratio maps were performed by careful visual inspection of previously tagged images by the same trained researcher, followed by inspection of lateral and medial snapshots of all T1w/T2w ratio maps. A few subjects were excluded at this final step, due to poor T1w/T2w ratio maps (n=34), ensuing a final sample of 621 subjects.

*Cognitive assessment*

During the cognitive flexibility task (Dimensional Change Card Sort Test), subjects had to match a picture to shape or color based targets, and the sum score was based on accuracy and reaction time incorporated by a two-vector method. During the inhibition and attention task (Flanker Inhibitory Control and Attention Test), subjects had to assess symbol orientation while ignoring surrounding and at times incongruent stimuli. While the youngest children were presented only with fish, subjects aged nine and above were additionally presented with smaller arrows. The inhibition sum score was based on congruent and incongruent trials, while the attention sum score was based on congruent trials only, both derived from a two-vector model. The episodic memory task (Picture Sequence Memory Test) involved recalling sequences of pictured objects and activities in the order they were presented. The length of the sequences presented depended on subject age. The sum score was based on the total number of pairs (two following pictures) placed in correct sequence order. During the processing speed task (Pattern Comparison Processing Speed Test) subjects were presented with pairs of pictures, and the sum score was based on total correct assessments of whether the pictures were identical or not. While subjects aged 8 and above pressed “yes” or “no” buttons, younger subjects correspondingly pressed a smiley or frowny face. During the reading test (Oral Reading Recognition Test) a word or letter was presented for the subject to read aloud. Letters and other multiple-choice items were presented for “pre-readers” and subjects with low literacy levels. The sum score was based on the total number of correct aloud readings of words and letters. The working memory task (List Sorting Working Memory Test) was a two-condition assignment where subjects were instructed to remember a series of presented objects and repeat them in size order. The sum score was based on total correctly remembered objects repeated in size order across conditions. The vocabulary task (Picture Vocabulary Test) consisted of responding to which picture most closely represented a word. The sum score was an item response theory theta conversion (Akshoomoff et al., 2014).

*Possible consequences of removing the t2pial flag during recon-all*

We did not find associations between T1w/T2w ratio and age in sub regions of medial occipital lobe and minor medial parietal and frontal regions, which was unexpected. This could be explained by our removal of the t2pial flag during the HCP pipeline, which when employed additionally uses the T2 image for pial surface estimation. As described, this removal was done to correct for sizable hemispheric differences induced in both thickness and T1w/T2w ratio values, possibly due to sequence or resolution differences as compared to the HCP standard. After careful visual inspection for the T1w and T2w image separately, we suspect that pial estimation within these regions was hampered by inclusion of dura in the T1w image alone, resulting in underestimation of T1w/T2w ratio values.

**Supporting Tables**

|  | Scanner 1 | Scanner 2 | Scanner 3 | Scanner 4 | Scanner 5 | Scanner 6 | Scanner 7 |
| --- | --- | --- | --- | --- | --- | --- | --- |
| N | 104 | 104 | 66 | 125 | 109 | 14 | 99 |
| Age (years) | 3.8-21.0 (mean= 14.9, SD= 4.0) | 3.2-21.0 (mean= 12.8, SD = 5.1 | 3.2-21.0 (mean= 14.4, SD = 5.4 | 3.2-20.3 (mean= 8.6, SD = 3.5 | 4.2-21.0 (mean= 10.9, SD = 3.6 | 8.8-20.8 (mean= 14.5, SD = 3.8 | 6.1-20.7 (mean= 14.2, SD = 4.0 |
| Sex | Boys= 61, Girls= 43 | Boys= 51, Girls =53 | Boys= 32, Girls =34 | Boys= 61, Girls =64 | Boys= 58, Girls =51 | Boys= 6, Girls =8 | Boys= 49, Girls =50 |
| Manufacturer | GE medical systems (signa HDx) | Siemens (TrioTim) | Siemens (TrioTim) | GE medical systems (Discovery MR 750) | Siemens (TrioTim) | Siemens  (TrioTim) | Siemens  (TrioTim) |

Supporting Table 1. Sample demographics per scanner. The table shows basic sample demographics for subjects within each scanner employed in the current study.

**Supporting Figures**

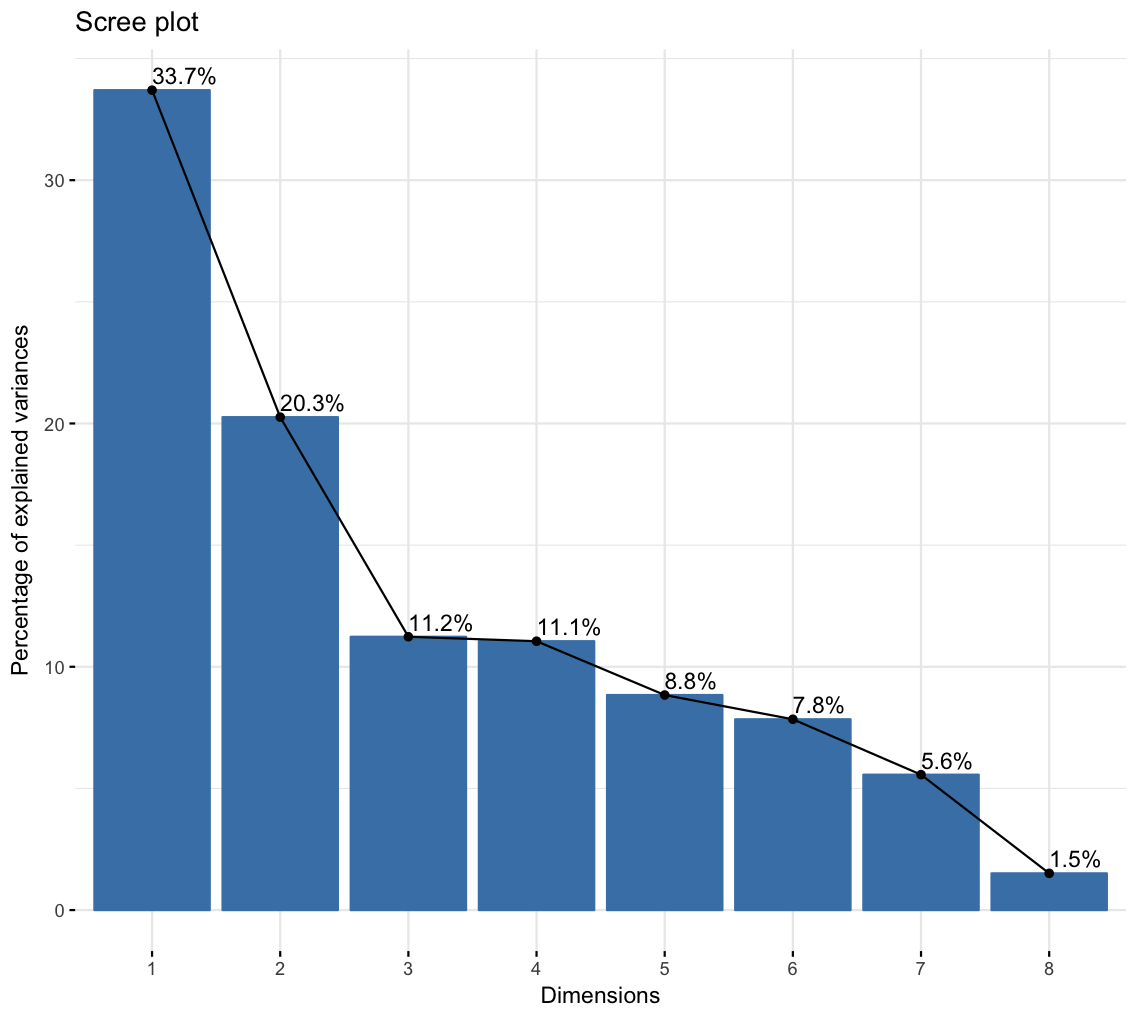

Supporting Figure 1: The percentage explained variance for the principal components. The figure shows a bar plot with the percentage explained variance for each of the 8 principal components retrieved from the principal component analysis on the cognitive sum scores.

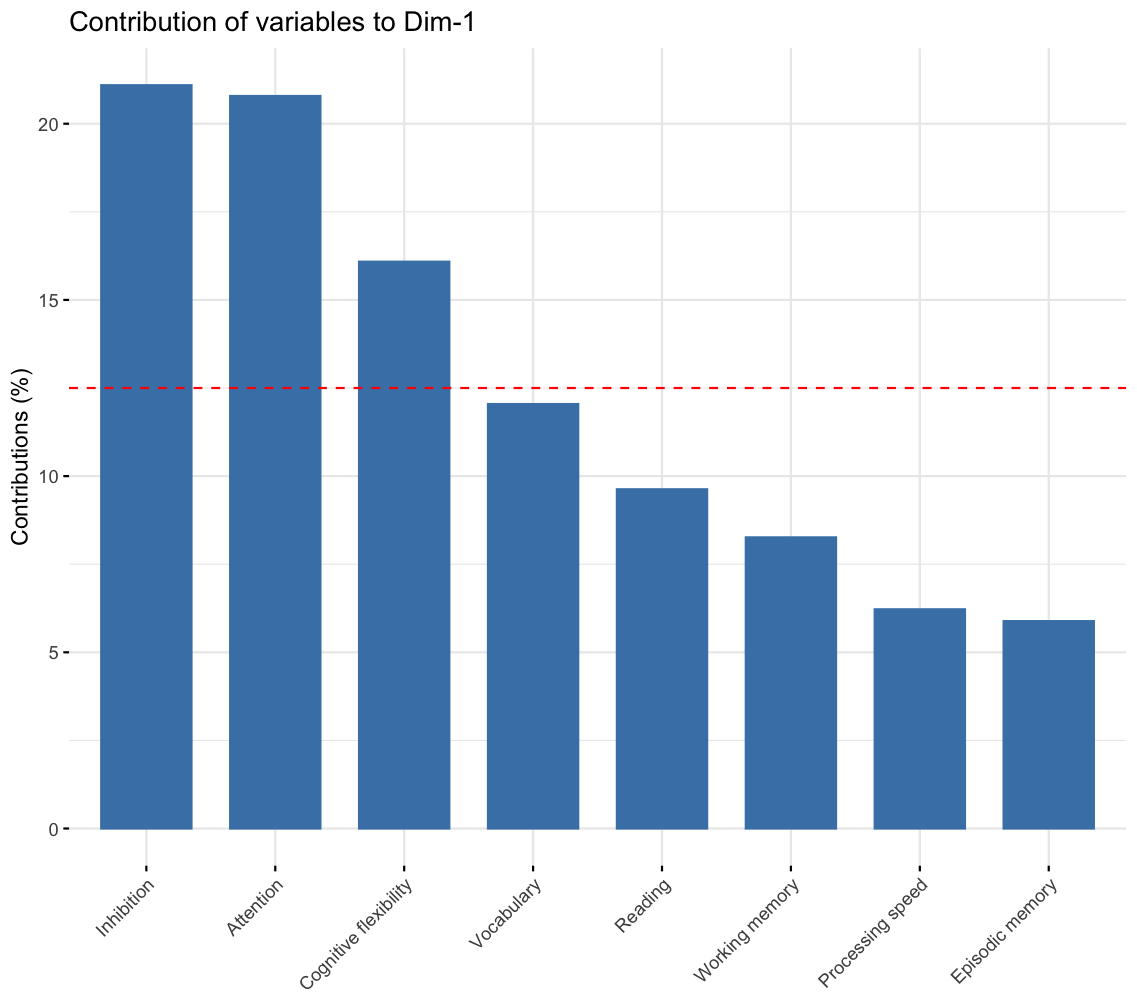

Supporting Figure 2. The percentage contribution of each cognitive sum score to principal component one. The figure shows a bar plot with the percentage contribution for each of the 8 sum scores to the first principal component which was extracted as a measure of general cognitive ability. The dashed red line is a reference line corresponding to the expected value if contributions were uniform.

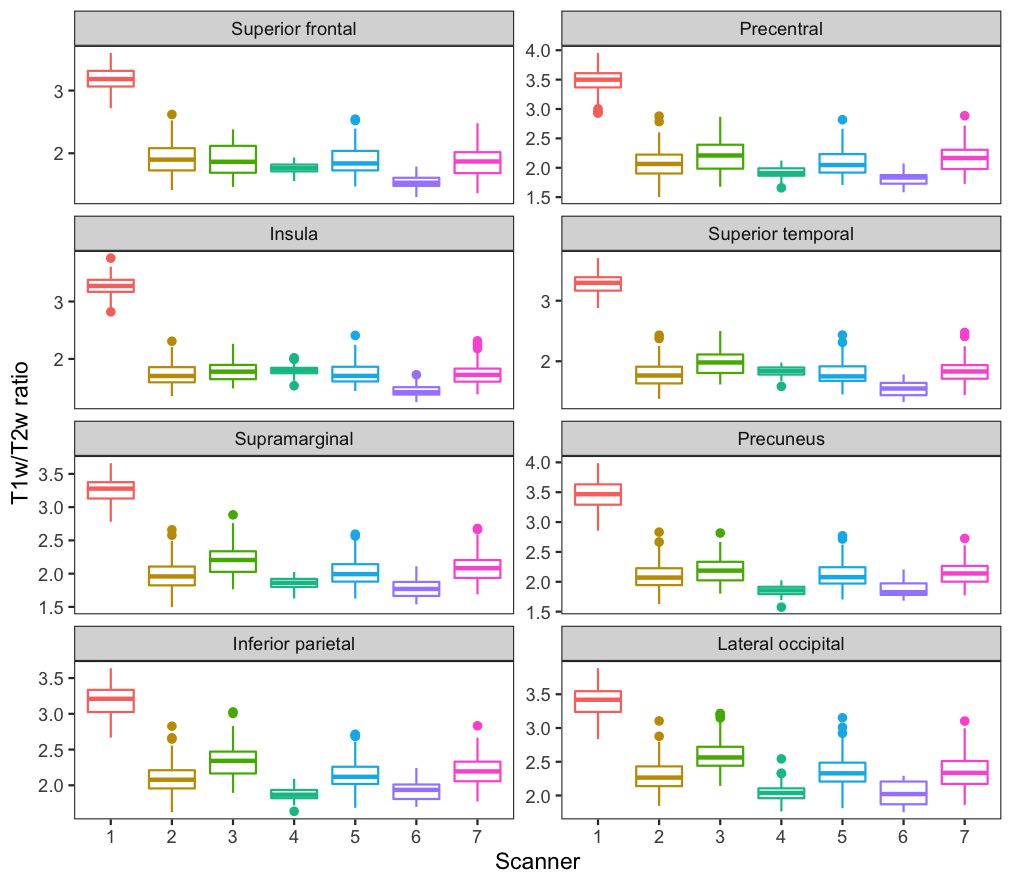

Supporting Figure 3. Regional box plots of T1w/T2w ratio intensity spread across scanners. The plots show T1w/T2w ratio intensity spread within a selection of regions from different cortical lobes for each scanner.

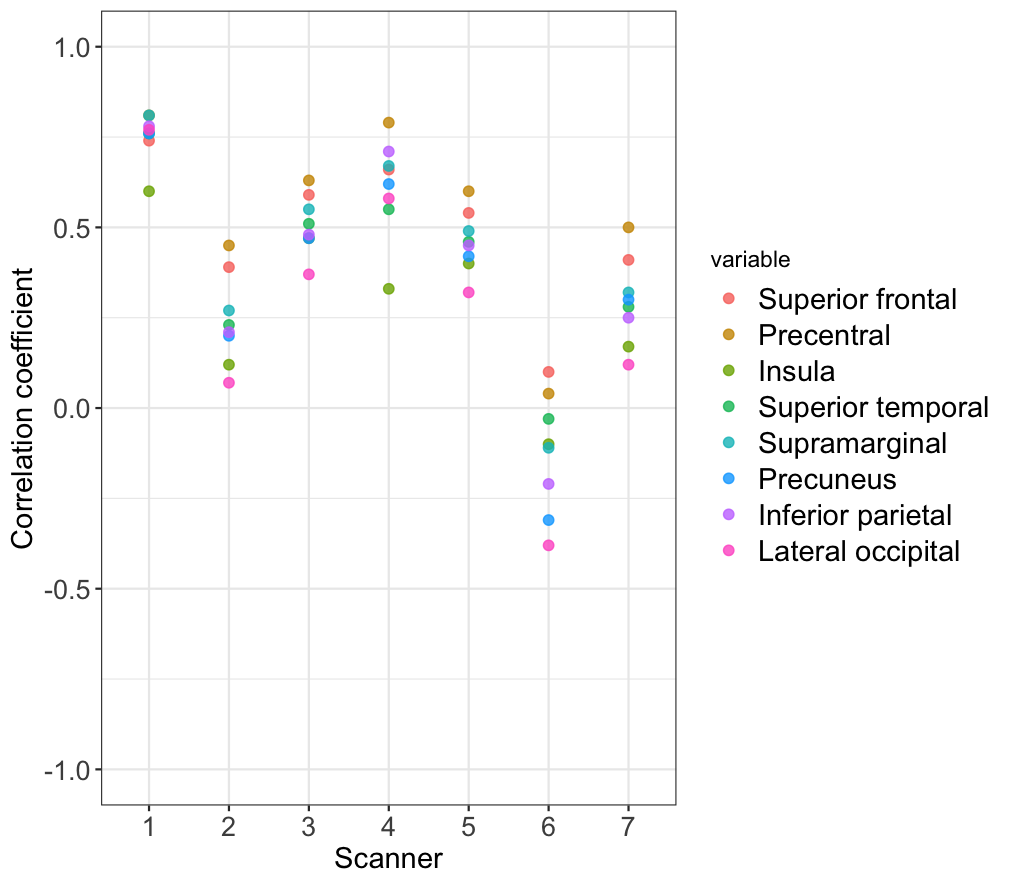

Supporting Figure 4. Scatter plot of correlations between regional T1w/T2w ratio and age. The figure shows Pearson’s correlation coefficients for age and T1w/T2w ratio within a selection of regions from different cortical lobes for each scanner.

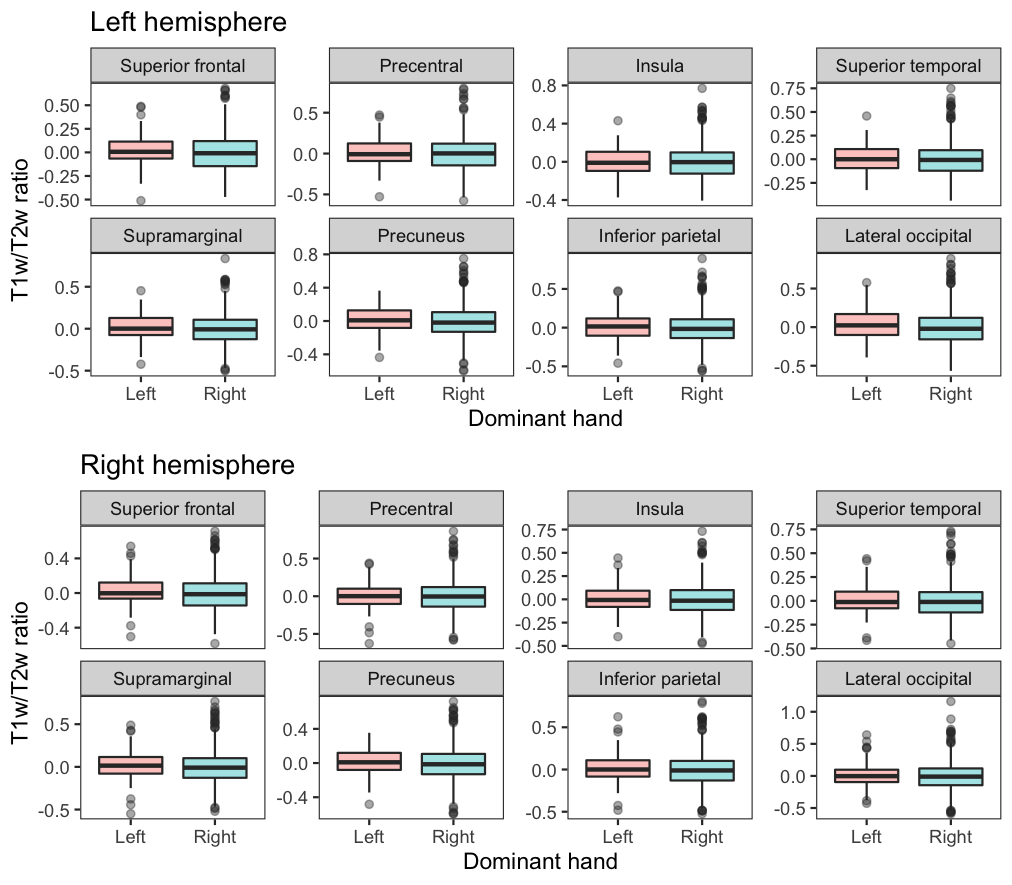

Supporting Figure 5. Box plots of the spread of T1w/T2w ratio within individuals with left- and right hand dominance. The figure shows box plots of mean T1w/T2w ratio within a selection of regions from different cortical lobes for both the left hand right hemisphere. T1w/T2w ratio is residualized for scanner.
